# Measurement of curvature and twist of microtubule bundles in the mitotic spindle

**DOI:** 10.1101/2021.01.02.425081

**Authors:** Arian Ivec, Monika Trupinić, Iva M. Tolić, Nenad Pavin

**Author notes:** Correspondence (A.I.), (N.P.).

## Abstract

The highly ordered spatial organization of microtubule bundles in the mitotic spindle is crucial for its proper functioning. The recent discovery of twisted shapes of microtubule bundles and spindle chirality suggests that the bundles extend along curved paths in three dimensions, rather than being confined to a plane. This in turn implies that rotational forces exist in the spindle in addition to the widely studied linear forces. However, studies of spindle architecture and forces are impeded by a lack of a robust method for the geometric quantification of microtubule bundles in the spindle. In this paper, we describe a simple method for measuring and evaluating the shapes of microtubule bundles, by characterizing them in terms of their curvature and twist. By using confocal microscopy, we obtain three-dimensional images of spindles, which allow us to trace the entire microtubule bundles. For each traced bundle, we first fit a plane, and then fit a circle lying in that plane. With this easily reproducible method, we extract the curvature and twist, which represent the geometric information characteristic for each bundle. As the bundle shapes reflect the forces within them, this method is valuable for the understanding of forces that act on chromosomes during mitosis.

## Introduction

Equal division of the genetic material into two newly formed daughter cells is performed by the mitotic spindle, a complex micro-structure that consists of two poles, microtubule bundles extending between the poles, and a large number of associated proteins (1–3). The spindle is a mechanical assembly that generates and regulates forces required for the segregation of chromosomes. The mechanical properties of the spindle arise from the mechanical properties of its basic building blocks, the microtubule bundles. Microtubules are thin elastic filaments that generate and balance the forces acting on chromosomes, which arise from the activity of motor proteins as well as from polymerization and depolymerization of the microtubule bundles (4).

It is feasible to directly measure the forces exerted on microtubule bundles (5), however it is rather challenging because of the small scales involved. To complement such experiments, it is possible to measure the forces indirectly by inferring them from the shape of the microtubule bundle (6–8). This approach is based on the fact that microtubules are inherently straight, but can obtain different shapes depending on the forces acting upon them. This approach was used to quantify the bending rigidity of single microtubules (8) and microtubule polymerization forces (7).

A similar approach for the quantification of forces based on shapes can be used on microtubule bundles in the spindle. The shapes of individual microtubule bundles give the entire spindle its characteristic shape, such as mitotic spindles in human cells. Similar spindle shapes can be found in most metazoans (9). Interestingly, even in spindles without centrosomes, e.g. in some protozoan organisms such as amoebas, a similar spindle shape is present (10). The same is the case for plant spindles (11). In some lower eukaryotes, e.g. yeasts, this type of spindle shape is absent as their spindles consist of a single straight microtubule bundle (12). Given that the spindle shape reflects the forces within it, accurate measurement and characterization of the shapes of microtubule bundles is highly important for the understanding of forces that act on chromosomes during mitosis.

We have recently shown that the spindle in human cells is a chiral object, as bundles follow a left-handed helical path (13). The chirality of the spindle is best visualized by looking at the spindle end-on, i.e., along the pole-to-pole axis, to observe the three-dimensional shapes with a helical twist. This view allows visualization of the microtubule bundles as flower petals (Fig. 1). By following the bundles in the direction towards the observer, the petals will rotate clockwise if the bundles follow a left-handed helical path. Vice versa, the petals will rotate counter-clockwise if the bundles follow a right-handed helical path.

**FIGURE 1.**
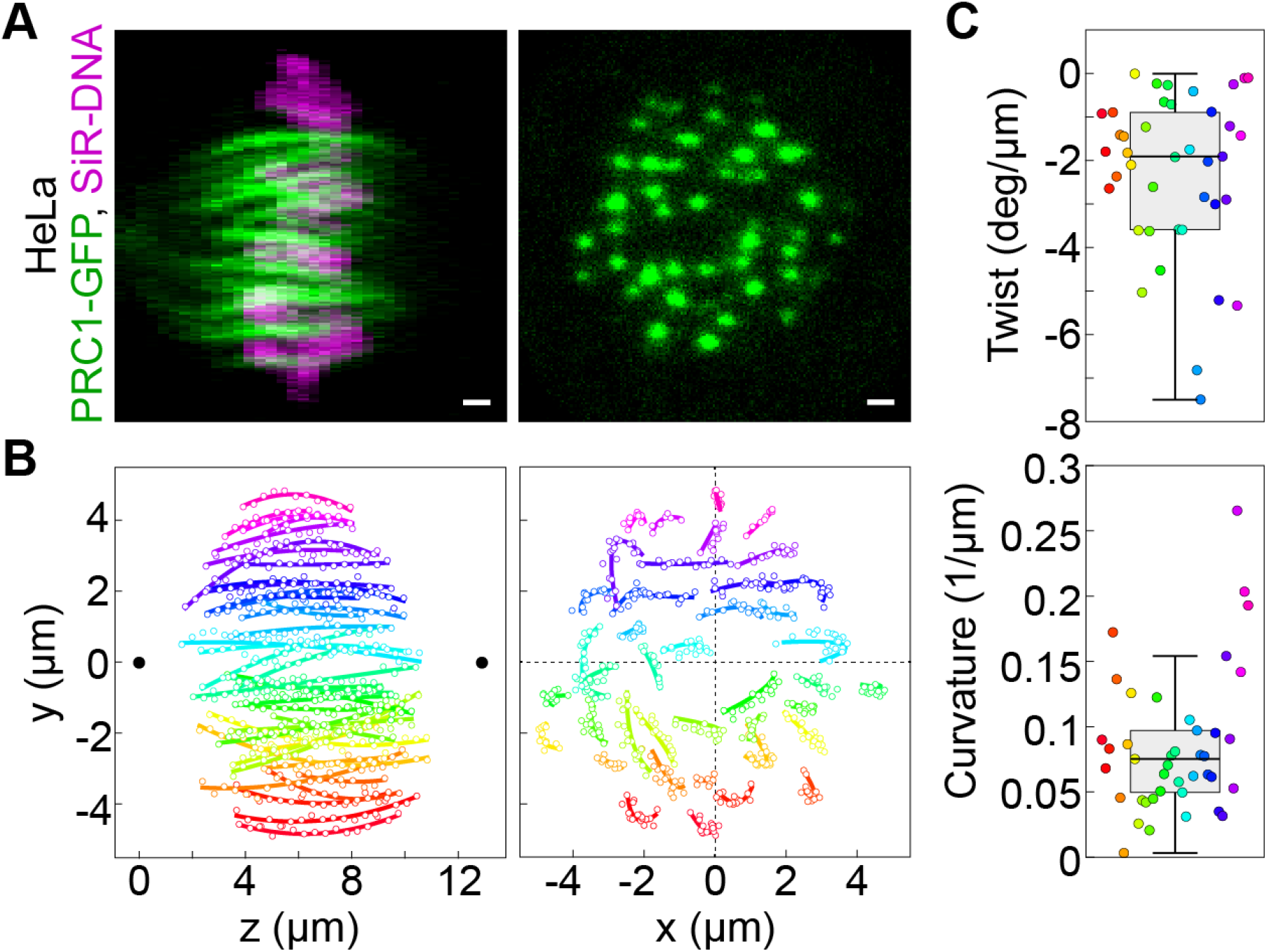
End-on view and side view of human mitotic spindle. (A) Confocal images of the metaphase spindle in a live HeLa cell expressing PRC1-GFP labelled with SiR-DNA dye shown in the end-on view (left, single z plane, only PRC1-GFP is shown) and side view (right, maximum intensity projection of the entire spindle). Scale bars, 1 μm. (B) Tracked microtubule bundles from the spindle in A shown in the end-on view (left) and side view (right). Each bundle is represented by a different color, thin circles mark the manually traced points along the bundle, and thick lines show circular arcs of the fitted circles. The centers of the centrosomes are represented as black dots in the side view. (C) Box plots (median and IQR=1.5) of the curvature and twist of each bundle are shown. The live HeLa cell shows a strong right (clockwise) twist.

The reason for this chirality may be the action of the motor proteins that exert rotational forces on the microtubules, such as kinesin-5 (Kif11/Eg5) (14), whose inhibition leads to the abolishment of left-handed twist (13). Motor proteins generate these forces by ‘walking’ along microtubules and performing a side-step while switching microtubule protofilaments (15–17).

In this paper, we develop a method for the analysis and measurement of the geometrical properties of microtubule bundles within the spindle. To extract the information about the shape of a microtubule bundle from noisy experimental data, we introduce a robust approach in which we consider the microtubule bundle as a part of a circular arc. Microtubule bundles can be approximated by thin curved lines that extend in three-dimensional space. This description allows us to faithfully represent the microtubule bundle and extract the relevant geometrical information from it, but it is also simple enough to be done systematically on a wide variety of microtubule bundles. Alongside the description of the method, we present a step-by-step application to a chosen microtubule bundle, to ease the understanding of the method.

## MICROTUBULE BUNDLE APPROXIMATED BY A CIRCLE

### Image analysis and data tracking

Individual microtubule bundles need to be tracked in order to acquire their *x, y, z* coordinates in each z-plane of the entire z-stack. Examples of such microtubule tracking can be found in (13). Each spindle has two poles positioned along the pole-to-pole axis, along with *N* microtubule bundles denoted by index *i* = 1,..,*N*. The *i*-th bundle is represented by set of *n_i_*, tracked points T_*ij*_ = (*x_j_,y_j_,Z_j_*). where *j* = 1,..,*n_i_* is the index of individual points (Fig. 2 *A*). Each bundle is tracked through all *z*-planes in the direction from left centrosome towards the right centrosome (the left centrosome represents the bottom z-plane, while the right centrosome represents the highest tracked z-plane in the stack). The positions of the centrosomes are the starting and end points of the spindle, so we include this information by extending the coordinates of each single bundle with the centrosome coordinates, with the left centrosome as the starting data point, T_i0_, and the right centrosome as the ending data point, T_ini +1_ (Supplementary Table 1), and thus coordinates of the *i*-th bundle are indexed *j* = 0,.., *n_i_* + 1. The *z*-plane refers to the imaging plane, which we convert to its corresponding *z*-coordinate by multiplying with the distance between successive planes set during image acquisition and by a factor of 0.81 to correct for the refractive index mismatch (13). In the example case (Fig. 1) the distance between z-planes is equal to 405 nm after correction, while other details of sample preparation are provided later in the text.

**FIGURE 2.**
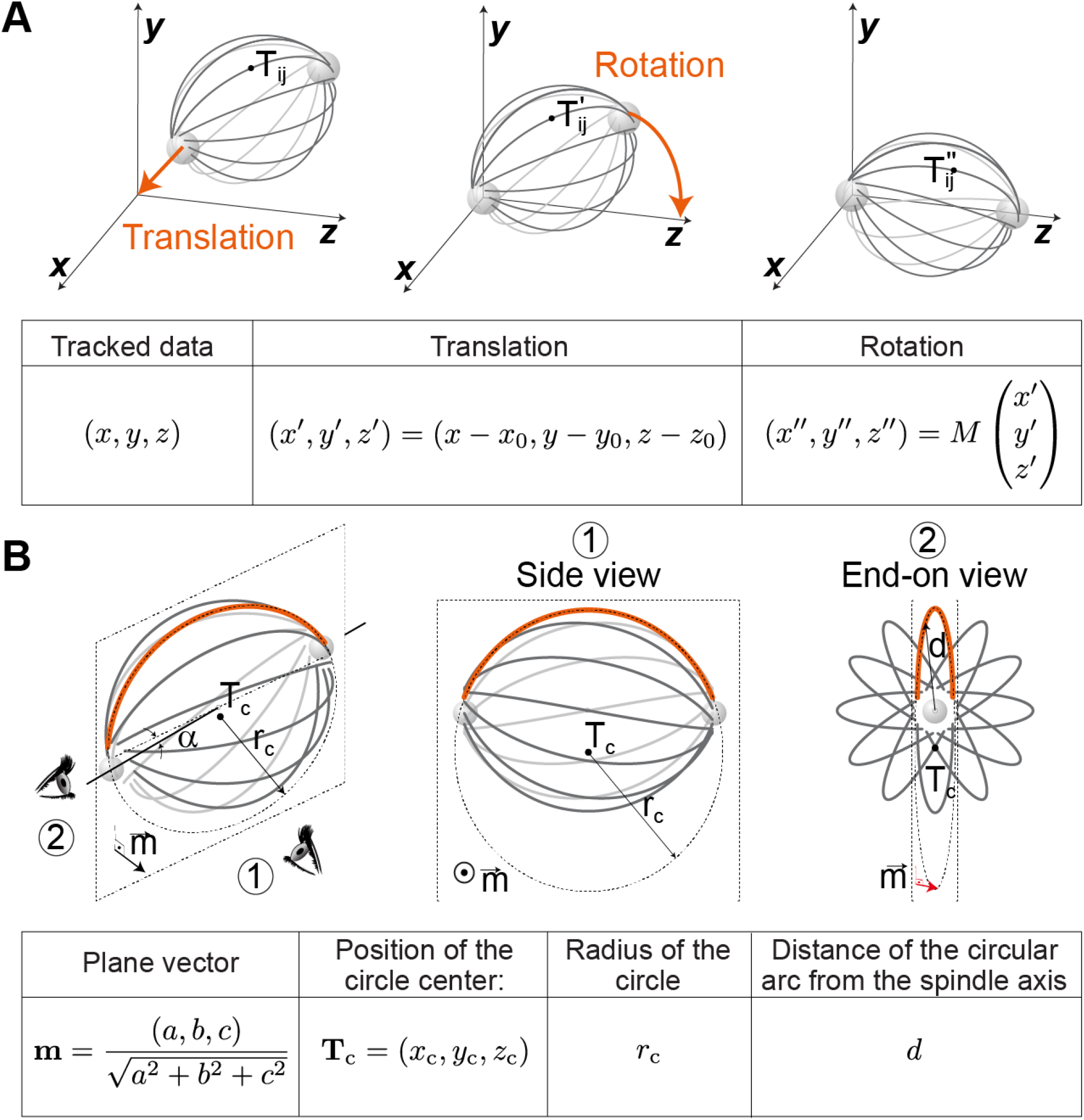
Overview of method (*A*) The spindle, along with the centrosomes and the marked traced bundle point T_ij_, is positioned at an arbitrary angle and distance from the origin of the coordinate system (left). The spindle is translated so that the left centrosome is located at the origin of the coordinate system (middle). The spindle is rotated so that the pole-to-pole axis, along with the right centrosome, aligns with the z-axis of the coordinate system (right). *(B)* A view of the spindle from an arbitrary angle (left) where eye signs mark the view angle for the side view (1) and the end-on view (2), which are shown in the middle and on the right, respectively. A microtubule bundle (orange curved line) is fitted by a circle of radius *r_c_*. The angle between the central spindle axis (solid line) and the plane in which the fitted circle lies (dashed parallelogram) is denoted. The parameters used to calculate the twist and curvature are named at the bottom of the scheme.

### Choosing a coordinate system

During imaging, spindles have an arbitrary location and orientation with respect to the laboratory coordinate system. To make tracks of microtubule bundles suitable for analysis, we transform the laboratory coordinate system so that the left centrosome is positioned at the origin of the new coordinate system and the right centrosome is positioned on the z-axis (Fig. 2 *A*), which we term the spindle coordinate system. The spindle coordinate system is obtained by two transformations: (i) translation 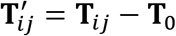, where 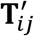 is transformed coordinate and T_0_ is position of the left centrosome (*T_0_* is given by the first row of Supplementary Table 1), and (ii) subsequent rotation 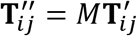, where 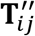 are coordinates in the spindle coordinate system and *M* is the rotation matrix which aligns the pole-to-pole axis with the *z*-axis of the spindle coordinate system and the unit vector 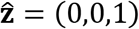. The rotation matrix is a textbook problem and it can be calculated, e.g., as the Rodrigues rotation (18) matrix. Finally, it is convenient to parameterize points by using cylindrical coordinates, 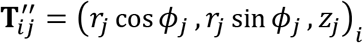, where *r_j_*, *ϕ_j_*, and *Z_j_* are, respectively, the radius, azimuth and axial position.

### Fitting a circular arc to the microtubule bundle shapes

To characterize complex three-dimensional shapes of microtubule bundles from noisy experimental data requires a robust approach. In our method, curvature and twist, which measure the extent the bundles extend along curved paths in three dimensions, are the geometrical quantities that represent the information about the bundle shapes. To obtain these quantities from the experimental traces, we choose to fit a circular arc extending through three dimensions to these data. Fitting the simple shape of a circular arc is a straightforward approach to extract the most important geometrical parameters from the data obtained from confocal microscopy, because the low number of data points we have for microtubule bundles makes the techniques of signal processing (19, 20) unapplicable. The more accurate data of bundle shapes obtained from super-resolution or electron microscopy might motivate more complex fitting shapes, which would make it possible to obtain twist and curvature directly (21).

To fit a circular arc in an easily reproducible way, we first fit a plane, and then fit a circle which lies in this plane (Fig. 2 *B*). Because we focus on one microtubule bundle only, we omit the bundle index *i* in the rest of this section.

In the first step, we fit the traced data to the best-fitting least squares distance plane *ax* + *by* + *cz* + *d* = 0, where are *a, b, c*, and *d* are the parameters of the general form equation of the plane, which we term the bundle plane. We solve the total least squares problem by using the singular value decomposition method (22). The normal unit vector of the bundle plane is given by 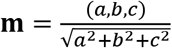

In the second step, we fit a circle to the data by choosing only from those circles that lie in the bundle plane. We calculate the projection of the traced bundle points onto the bundle plane and fit a circular arc to them. For a given set of points in the plane, this method of fitting a circle provides a unique closed form solution, which minimizes the corresponding orthogonal distance. The fitting parameters are the radius of the circle, *R_c_*, and the position of the circle center, T_c_ = (*x_c_, y_c_, z_c_*). These parameters, together with the normal vector of the bundle plane, determine the geometry of our traced bundle.

We note that fitting a circle with standard methods (23) in most cases produces acceptable results, with the outliers straight bundles, who require circles of larger radii. We thus use the HyperLS (24) algorithm to both easily handle such cases, and to benefit from a closed form solution.

### Calculation of the curvature and twist from the fitting parameters

Based on the fitting parameters, we can infer the curvature and twist of the microtubule bundle. The curvature of the bundle can be directly calculated from the radius of the fitted circle,

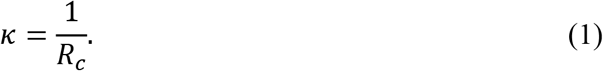

The twist, however, cannot be calculated in a straightforward manner. We introduce the twist value, *ω*, as a change of the azimuthal angle with respect to the axial position

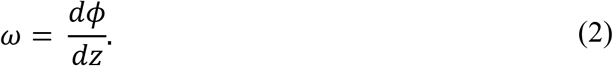

Please note that this value corresponds to the reciprocal value of helical pitch multiplied by 2π. Because helical trace can be approximated by a circular arc for small value of dφ, the right-hand side of Eq. (2) can be calculated as *d*φ/*dz* = (cotα)/*r*. Here, the circular arc radius corresponds to the radial coordinate *r* in the spindle coordinate system, and *α* denotes the angle between the plane of the circular arc and z-axis. We calculate this angle from the scalar product cos 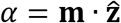. For our case of discrete tracked bundle data point, we average the radius over all traced points 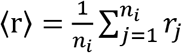, and thus Eq. (2) can be written as

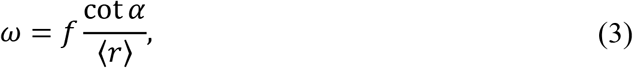

where *f* is a dimensionless corrective factor, which arises because of the approximative approach of the method due to fitting a circular arc to the bundle segment. The corrective factor depends on the geometry of the bundle, but for short bundle segments, significantly shorter than the spindle length, one can use *f* = 1.

### Detailed worked example - synthetic spindle

To demonstrate the workings of our method, we provide a detailed worked example on a made-up mitotic spindle in a spindle coordinate system, which mimics the one shown in the schematic in Fig. 2 *B*, but also includes noise to make it closer to experimental data. We construct a synthetic spindle as a series of mathematically defined curves, which are evenly distributed around the *z*-axis. To do this, we first define a bundle as a twisted circular arc:

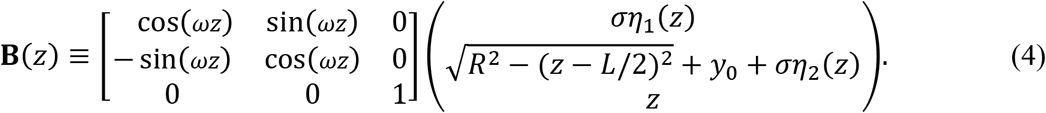

The first term is a matrix which twists the bundle by *ω* around the *y* — axis. The second term is a vector that defines the bundle as a circular arc with added noise, where *R* is radius of the circular arc, *L* is the total length of the spindle, equal to *L* = 2*R*, and constant 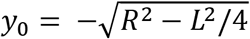 is chosen such that the circular arc extends from one pole to the other. The two components of noise, η_1,2_(z), are Gaussian white noise, which obey 〈η_β_(z)η_γ_(z‘)〉 = δ(z — z’)δ_β,γ_, with δ(z — z’) being the Dirac *δ* function and δ_β,γ_ is the Kronecker *δ* function.

To obtain a spindle, we evenly arrange the *N* bundles around the pole-to-pole axis, so that the *i*-th bundle, T_i_, is given by

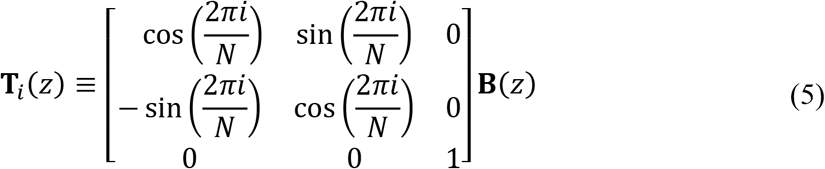

The first term represents a matrix which rotates the *i*-th bundle, in order to obtain a spatial configuration of bundles distributed around the *z*-axis.

To mimic discrete imaging planes, we choose to assign to the *z*-coordinates discrete values *Z_j_* = *L*_0_ + *jΔL*, where *j* = 1,…,*n_i_*. Here, *L*_0_ + *ΔL* and *L*_0_ + *n_i_ΔL* denote the starting point and ending *z*-coordinates of the synthetic bundle segments. Finally, positions of spindle poles are given by

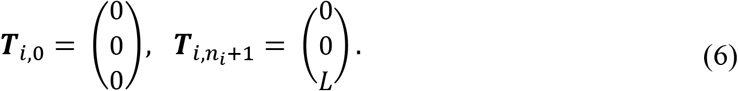

Our method will measure the defined twist of this curve *ω*, as well as estimate its curvature.

### Fitting circular arcs to the bundles of synthetic spindle

We fit a circular arc to the synthetic bundles by using the approach described in the section Fitting a circular arc to the microtubule bundle shapes. In the first step, we obtain parameters of the bundle plane, *a, b, c*, and *d*. In the second step, we fit the circular arc which lies in the bundle plane and the corresponding radius R_c_.

To test our method, we apply the method to four synthetic spindles shown in Fig. 3. We have chosen two spindles, the first with short bundle segments and the second with long bundle segments, each with and without noise. In the case with short bundle segments the twist and curvature obtained from the method closely matches parameters which define the synthetic spindle, both with and without noise (Fig. 3 *A, B*). This agreement is expected because the fitting curve closely follows the synthetic bundle segments in the case without noise. In the case with long bundle segments, the obtained twist is slightly smaller than the defined one and the difference between the fitted curve and the synthetic bundle segments becomes visible (Fig. 3 *C, D*).

**FIGURE 3.**
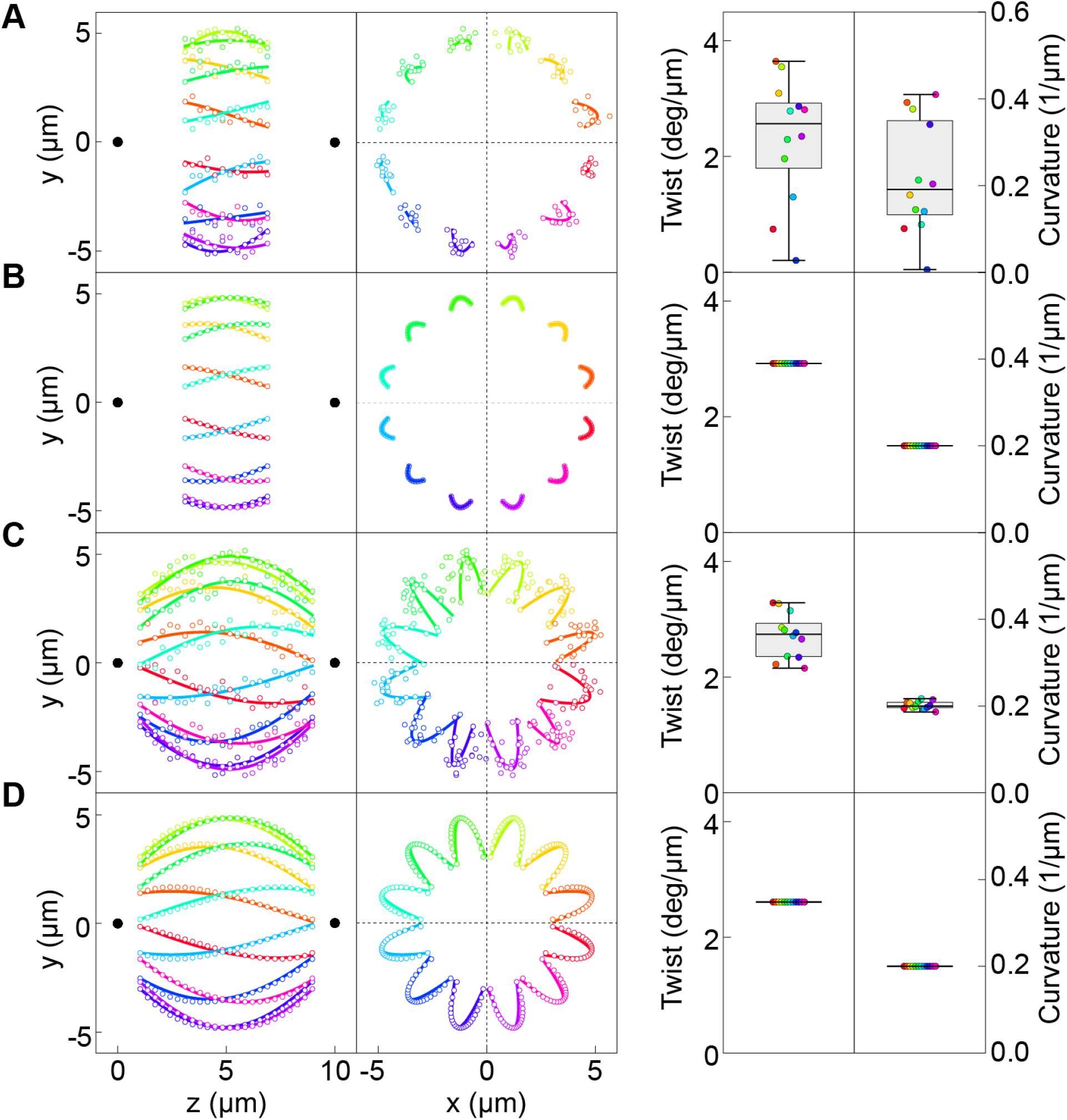
Application of method to four synthetic spindles. We show synthetic spindles (colored circles) together with poles (black points), fits to them (colored lines) in the side view (left) and in the end-on view (middle), along with corresponding values of twist and curvature (right). The first two spindle have short bundle segments, in which 40% of central bundles were calculated, both with noise (A) and without noise (B). The last two spindles have long bundle segments, in which 80% of central bundles were calculated, with noise (C) and without noise (D). Short segments are comprised of *n* = 12 points, with noise *σ* = 0.25 in (A) and *σ* = 0 in (B). Long segments are comprised of *n* = 24 points, with noise *σ* = 0.25 in (C) and *σ* = 0 in (D). For all spindles, the values of the other parameters are *ΔL* = 0.33 *μm*, *ω* = 3 *deg/μm* and *R* = 5. Twist and curvature are shown with individual points and box plots (median and IQR=1.5).

Because in the case of long bundle segments the twist we obtain from our method underestimates the exact value, we explored how the discrepancy changes with the length of the bundle segment. We calculate the twist for bundle segment length ranging from 0-10 μm for a spindle with pole-to-pole distance of 10 μm and plot the value of the twist obtained from our method divided by the exact value, shown in Fig. 4. We also tested the dependence of the discrepancy for values of parameters *B* and *ω*. For *B* = 1 μm and 5 μm, which correspond to inner and outmost bundles in HeLa cells, our numerical calculations show that the difference is small and that it does not exceed 1‰. Similarly, for *ω* = 1 deg/μm and 3 deg/μm, values typical for HeLa cells, is also small and does not exceed 1‰. The value of the twist obtained by our method systematically underestimates the exact value for longer bundle segments, whereas the other parameters which define the shape of the synthetic spindle change the value negligibly.

To obtain, by our method, a value of the twist as close as possible to the exact value, we need to calculate the corrective factor in Eq. (3). Because this factor predominately depends on the bundle segment length, we calculate it as a function of the normalized bundle segment length, *t* = *n* Δ*L*/*L*. Approximatively, this function is given by

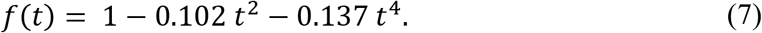

Because the synthetic spindle is similar to spindles found in HeLa cells, this phenomenological function can be used for all bundles in HeLa spindles, and also for spindles that have shapes similar to HeLa cells.

### Comparison with previous results

In the study Novak et al. (13) twist was calculated for short bundle segments in HeLa and U2OS cells by using Eq. 2 for finite segments Δz. Here, we calculate the values of the twist for the same bundle segments by using the method developed in this paper. In the example in Fig. 1 we obtain value of twist to be *ω* = 2.3 ± 0.3 deg/μm, which is consistent with value of twist resulting from the method in Novak et al. (13), by which the twist for this cell would be equal to *ω* = 2.3 ± 0.4 deg/μm. Though the obtained values are similar for short bundle segments, the method developed here is robust and can be applied to a greater variety of microtubule bundles, including those with longer bundle segments and bundles closer to the pole-to-pole axis.

### What we learn from the curvature and the twist

Curvature and twist provide geometrical information about the shape of the bundle. Based on these geometrical parameters, we can infer information about rotational forces, i.e., bending and twisting moments. Curvature can provide an estimate of the bending moment acting upon the bundle

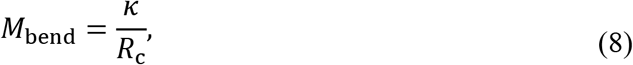

where R_c_ is the radius of curvature measured for a microtubule bundle and *κ* is its flexural rigidity.

Twist characterizes to which extent bundles rotate around the spindle pole-to-pole axis. Intuitively, one can expect that twist is related to twisting moment within these bundles. This is indeed the case for spindles described by model from (13), in which microtubule bundles, that are intrinsically straight, extend radially from spindle poles. In this case, we can obtain an estimate for the twisting moment acting upon the microtubule bundle

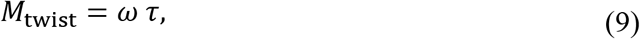

where *ω* is the measured twist and *τ* is the torsional rigidity of the microtubule bundle (25).

## EXPERIMENTAL METHODS

### Cell lines and microtubule visualization

Twist can be measured in every cell line that has labelled microtubule bundles. This label can be a fluorescent protein tag on a microtubule bundle (e.g. on tubulin) or on proteins that are associated with microtubule bundles in a way that they cover most of the length of the bundle (e.g. PRC1, see Fig. 1). Tags can be inserted into the cells on a plasmid by transfection or endogenously expressed after CRISPR/Cas manipulation. Dyes, such as SiR-tubulin (26), can also be added to visualize microtubule bundles. It is possible to measure twist both in live and fixed cells. In fixed cells it is important to perform an appropriate fixation method. Fixation with methanol can often cause spindles to shrink in z-direction, which yields measurements that are not relevant for live cells. Spindles in fixed cells need to closely resemble spindles in live cells where spindle length and width need to be the same. In fixed cells, fluorescently labelled antibodies can also be used for tubulin visualization. Examples of imaging both live and fixed cells for purpose of measuring twist can be found in Novak et al. (13).

### Confocal microscopy

To measure twist, whether in live or fixed cells, imaging of the entire spindle needs to be performed. This means that the imaged z-stack needs to be big enough to encompass the spindle from the bottom of the dish to the top of the spindle. Spindles that are oriented horizontally (spindle pole-to-pole axis is parallel with the imaging plane) or vertically (spindle pole-to-pole axis is perpendicular to the imaging plane) are the most appropriate for the analysis. In both cases it is important to capture the entire spindle. Also, it is important to note that the direction of the imaging needs to be from the glass bottom upwards. The bottom of the dish is usually easy to determine by surrounding cells in interphase that are attached there. The direction of the imaging is important for determining handedness of the twist (right-or left-handed twist). An example of microscope settings for the purpose of imaging spindles for measuring twist can be found in Novak et al. (13).

## CONCLUSION

The discovery that microtubule bundles in the mitotic spindle are twisted in a helical manner open an exciting area of research on the potential biological roles of spindle chirality and the mechanisms generating this curious type of asymmetry, which is why we developed a method to measure the twist and the curvature of microtubule bundles in order to characterize the shape of the spindle. The method allows for easy extraction of information about the relevant aspects of microtubule bundle geometry. By utilizing the characteristic shape of microtubule bundles in the spindle, it is possible to characterize them in a reproducible manner. This approach opens up new lines of studies, allowing for efficient mapping of the similarities and differences between shapes of spindles in various cell types and organisms. Because the spindle shapes reflect the forces within them, this method will be instrumental for the understanding of forces that act on chromosomes during cell division.

## ACKNOWLEDGMENTS

The authors thank Ina Poser and Tony Hyman (Max Planck Institute of Molecular Cell Biology and Genetics, Dresden, Germany) for HeLa-Kyoto BAC cell line stably expressing PRC1-GFP; Barbara Kokanović for the image of HeLa PRC1-GFP spindle used in Fig. 1 and all members of Pavin and Tolić groups for helpful discussions. Research in the Pavin and Tolić groups is supported by the European Research Council (ERC Synergy Grant, GA Number 855158, granted to I.M.T. and N.P.), Croatian Science Foundation (HRZZ, projects IP-2019-04-5967 granted to N.P., PZS-2019-02-7653 granted to I.M.T. and the work of doctoral students A.I. and M.T. has been fully supported/supported in part by the corresponding “Young researchers’ career development project - training of doctoral students”), the QuantiXLie Center of Excellence, a project co-financed by the Croatian Government and European Union through the European Regional Development Fund—the Competitiveness and Cohesion Operational Programme (Grant KK.01.1.1.01.0004). I.M.T. acknowledges earlier support from the ERC (Consolidator Grant, GA Number 647077).

## REFERENCES

1. McIntosh, J. R., M. I. Molodtsov, and F. I. Ataullakhanov. 2012. Biophysics of mitosis. Q Rev Biophys 45(2):147–207.

2. Pavin, N., and I. M. Tolic. 2016. Self-Organization and Forces in the Mitotic Spindle. Annu Rev Biophys 45:279–298.

3. Prosser, S. L., and L. Pelletier. 2017. Mitotic spindle assembly in animal cells: a fine balancing act. Nat Rev Mol Cell Biol 18(3):187–201.

4. Howard, J. 2001. Mechanics of Motor Proteins and the Cytoskeleton. Sinauer Associates, Publishers.

5. Nicklas, R. B. 1983. Measurements of the force produced by the mitotic spindle in anaphase. J Cell Biol 97(2):542–548.

6. Pavin, N., and I. M. Tolic. 2020. Mechanobiology of the Mitotic Spindle. Developmental Cell. DOI: 10.1016/j.devcel.2020.11.003

7. Dogterom, M., and B. Yurke. 1997. Measurement of the force-velocity relation for growing microtubules. Science 278(5339):856–860.

8. Gittes, F., B. Mickey, J. Nettleton, and J. Howard. 1993. Flexural rigidity of microtubules and actin filaments measured from thermal fluctuations in shape. The Journal of cell biology 120(4):923–934.

9. Crowder, M. E., M. Strzelecka, J. D. Wilbur, M. C. Good, G. von Dassow, and R. Heald. 2015. A comparative analysis of spindle morphometrics across metazoans. Current Biology 25(11):1542–1550.

10. Walsh, C. J. 2012. The structure of the mitotic spindle and nucleolus during mitosis in the amebo-flagellate Naegleria. PloS one 7(4):e34763.

11. Zhang, H., and R. K. Dawe. 2011. Mechanisms of plant spindle formation. Chromosome Res 19(3):335–344.

12. McCully, E. K., and C. F. Robinow. 1971. Mitosis in the fission yeast Schizosaccharomyces pombe: a comparative study with light and electron microscopy. Journal of cell science 9(2):475–507.

13. Novak, M., B. Polak, J. Simunic, Z. Boban, B. Kuzmic, A. W. Thomae, I. M. Tolic, and N. Pavin. 2018. The mitotic spindle is chiral due to torques within microtubule bundles. Nat Commun 9:63–70.

14. Yajima, J., K. Mizutani, and T. Nishizaka. 2008. A torque component present in mitotic kinesin Eg5 revealed by three-dimensional tracking. Nature structural & molecular biology 15(10):1119–1121.

15. Ramaiya, A., B. Roy, M. Bugiel, and E. Schäffer. 2017. Kinesin rotates unidirectionally and generates torque while walking on microtubules. Proceedings of the National Academy of Sciences 114(41):10894–10899.

16. Can, S., M. A. Dewitt, and A. Yildiz. 2014. Bidirectional helical motility of cytoplasmic dynein around microtubules. Elife 3:e03205.

17. Bormuth, V., B. Nitzsche, F. Ruhnow, A. Mitra, M. Storch, B. Rammner, J. Howard, and S. Diez. 2012. The highly processive kinesin-8, Kip3, switches microtubule protofilaments with a bias toward the left. Biophys J 103(1):L4–L6.

18. Rodrigues, O. 1840. Des lois geometriques qui regissent les desplacements d’un systeme solide dans l’espace et de la variation des coordonnees provenant de deplacements consideres independamment des causes qui peuvent les produire. J Mathematiques Pures Appliquees 5:380–440.

19. Friedrich, B. 2020. frenet_robust.zip (https://www.mathworks.com/matlabcentral/fileexchange/47885-frenet_robust-zip), MATLAB Central File Exchange. Retrieved October 21, 2020.

20. Friedrich, B. 2020. powersmooth (https://www.mathworks.com/matlabcentral/fileexchange/48799-powersmooth), MATLAB Central File Exchange. Retrieved October 25, 2020.

21. Jikeli, J. F., L. Alvarez, B. M. Friedrich, L. G. Wilson, R. Pascal, R. Colin, M. Pichlo, A. Rennhack, C. Brenker, and U. B. Kaupp. 2015. Sperm navigation along helical paths in 3D chemoattractant landscapes. Nat Commun 6:7985.

22. Golub, G. H., and C. Reinsch. 1971. Singular value decomposition and least squares solutions. Linear Algebra. Springer, pp. 134–151.

23. Taubin, G. 1991. Estimation of Planar Curves, Surfaces, and Nonplanar Space Curves Defined by Implicit Equations with Applications to Edge and Range Image Segmentation. IEEE Trans. Pattern Anal. Mach. Intell. 13(11):1115–1138.

24. Kanatani, K., and P. Rangarajan. 2011. Hyper least squares fitting of circles and ellipses. Comput Stat Data An 55(6):2197–2208.

25. Landau, L. D., E. M. Lifshitz, A. M. Kosevich, J. B. Sykes, L. P. Pitaevskii, and W. H. Reid. 1986. Theory of Elasticity: Volume 7. Elsevier Science.

26. Lukinavicius, G., L. Reymond, E. D’Este, A. Masharina, F. Gottfert, H. Ta, A. Guther, M. Fournier, S. Rizzo, H. Waldmann, C. Blaukopf, C. Sommer, D. W. Gerlich, H. D. Arndt, S. W. Hell, and K. Johnsson. 2014. Fluorogenic probes for live-cell imaging of the cytoskeleton. Nat Methods 11(7):731–733.

